# Improving coral oxidative stress assessments through compartment-specific lipid peroxidation measurements and increased methodological standardization

**DOI:** 10.64898/2026.07.08.737270

**Authors:** Sophia W. Mastorakos, Avery J. Kruger, Chloe Carbonne, Yvonne Sawall, Liza M. Roger

**Author notes:** Corresponding author: Liza M. Roger.

## Abstract

Lipid peroxidation (LPO) is widely used as a biomarker of oxidative stress in coral bleaching research, yet its measurement remains poorly standardized across the field. A systematic review of the coral LPO literature reveals substantial variation in methodological approaches, including tissue fraction analysis, lysis protocols, assay choice, and normalization metrics, confounding cross-study comparison and obscuring the biological interpretation of results. We experimentally investigate two key sources of variation: the use of bulk holobiont vs separated host and algal symbiont fractions, and the choice of normalization metric. To do so, we used *Montastraea cavernosa* (n = 6 colonies) exposed to ambient (28 °C), heat stress (30.5 °C), and heat stress + artificial upwelling (AU; heat stress intermitted by daily pulses of cooler water, 30.5/27.5 °C) conditions in a controlled mesocosm experiment. Using a TBARS-based MDA assay with a lysis buffer optimized for coral tissue, we measured LPO separately in coral host and algal symbiont fractions across four time points throughout the day. Host MDA remained stable across all treatments and time points, consistent with either sufficient antioxidant buffering capacity or thermal acclimation over the experimental period. Algal symbiont MDA, in contrast, exhibited pronounced diel and treatment-specific dynamics, and the two fractions’ responses were decoupled from one another. Normalizing MDA to coral surface area instead of total protein content produced largely consistent diel and treatment patterns, but the two metrics diverged at specific time points, indicating that normalization choice is not interchangeable and can itself affect interpretation. Together, our literature review and empirical results demonstrate that host and algal symbiont LPO dynamics are not comparable when aggregated and argue for host-symbiont fraction separation and consistent, explicitly reported normalization as minimum standards for interpretable and cross-comparable coral LPO measurement.

## 1. Introduction

Coral reefs have experienced unprecedented global decline over recent decades, with thermal bleaching events identified as a primary driver of coral mortality worldwide (Hughes et al 2018; Oliver et al 2018). Bleaching occurs when the symbiosis between corals and their dinoflagellate endosymbionts (family Symbiodiniaceae) breaks down, often fatally, if the relationship is not re-established (Davy et al., 2012; LaJeunesse, 2020). At the cellular level, oxidative stress has emerged as a central feature of the bleaching response, whereby overproduction of reactive oxygen species (ROS) and reactive nitrogen species (RNS) overwhelms the coral’s antioxidant defenses (Downs et al., 2002). A key downstream consequence of this oxidative burden is lipid peroxidation (LPO), the oxidative degradation of polyunsaturated fatty acids (PUFAs) in cellular membranes, which has the capacity to compromise membrane integrity, trigger apoptotic cascades, and propagate further oxidative injury (Juan et al 2021; Wang et al 2023).

Lipids are a diverse class of biological molecules that serve as fundamental building blocks of cellular architecture across all domains of life (Muro et al., 2014). They form the phospholipid bilayers of key cellular structures, including the nuclear envelope, endoplasmic reticulum, and plasma membrane, where they are essential for maintaining cellular homeostasis via transmembrane transport (Muro et al., 2014; Storck et al., 2018; Tang et al., 2021). Cellular lipid reserves are also essential to metabolic activity under normal and stressed conditions (Rodrigues and Grottoli 2007).

Due to the cnidarian’s obligate symbiosis with photosynthetic dinoflagellate algae, the coral holobiont presents a uniquely complex biochemical environment in which to study LPO compared to well-studied mammalian systems. Both partners contribute distinct lipid pools: the coral host is rich in storage lipids such as wax esters and triacylglycerols, while algal symbionts are characterized by high concentrations of PUFAs (Imbs, 2013; Imbs & Dembitsky, 2023; Yamashiro et al, 1999). The majority of coral research has focused on the holobiont, although the coral host and algal symbionts may express vastly different LPO dynamics.

Free radicals are important mediators in redox signaling, but their overaccumulation leads to oxidative damage and potentially to cell death (Murphy et al 2022). Both the coral host and algal symbiont generate free radicals endogenously via mitochondria and the endoplasmic reticulum activity, through a series of enzymatic and non-enzymatic reactions (Ayala et al 2014), with algal symbionts generating additional free radicals as a byproduct of photosynthetic electron transport (Lesser, 2006). Lipids are susceptible to free radical damage when overaccumulation occurs, initiating the chain reaction of oxidative membrane degradation that characterizes LPO (Yin et al 2011). Critically, since both partners contribute independent sources of oxidative stress, LPO signals measured in the holobiont reflect the combined and potentially confounded response of two biochemically distinct cell types, complicating interpretation relative to well-studied mammalian systems (Lesser, 2019; Szabo et al 2020; Downs et al 2002).

LPO is the process by which lipids are attacked by these free radicals, potentially causing severe membrane damage if not mitigated by intracellular antioxidant defense (Ayala et al 2014). While LPO can produce bioactive signaling molecules such as lipid peroxides, excessive LPO can result in lipotoxicity, compromising cellular lipid stability (Aschner et al 2021). LPO follows a well-characterized schematic of initiation, propagation, and termination (Supp. Info. Fig S1) (Yin et al 2011). During initiation, extra- or intracellular stress generates a lipid radical (Yin et al 2011). This lipid radical then reacts with molecular oxygen, shifting the reaction to the propagation phase and creating a peroxy radical (Atukeren, 2021; Gaschler et al 2017). The peroxy radical subsequently reacts with adjacent lipid compounds and PUFAs, progressively altering the structure and dynamics of lipid membranes until quenched by antioxidants or until apoptosis ensues (Gaschler et al 2017). LPO targeting PUFAs produces characteristic secondary products, including malondialdehyde (MDA) and 4-hydroxynonenal (4-HNE), that have been used as molecular indicators of membrane damage (Schneider et al., 2008; Zhang et al 2020).

Various assays have been developed to detect these molecules as proxies for LPO (Ayala et al 2014). Among these, MDA in particular reacts with thiobarbituric acid (TBA), producing a pink chromophore quantified using the Thiobarbituric Acid Reactive Substances (TBARS) assay (El-Aal, 2012). While the study of LPO in the context of coral thermal stress is increasing, methodologies are not standardized across the field (Szabo et al 2020; Sikorskaya & Imbs, 2020).

Previous studies of LPO in corals reveal substantial inconsistency in both absolute LPO data and reported trends across bleaching conditions (Table 1). Some variation across studies is expected given genuine biological differences in coral LPO responses across species, genotypes, colonies, and environment (Flores-Ramírez & Liñán-Cabello, 2007; Montalbetti et al., 2021; Pinheiro et al., 2026). However, substantial LPO variation has been reported even within species under comparable experimental conditions (Dorantes-Aranda et al., 2026; Montalbetti et al., 2021; Rädecker et al., 2021), raising concern that the methodological inconsistency is a contributing factor. When methodological and biological sources of variation cannot be disentangled, it becomes unclear whether reported differences in LPO reflect true physiological responses or experimental artifacts. Standardization of LPO methodology is therefore needed to enable meaningful cross-study comparison and to resolve whether observed variation is biologically informative.

**Table 1.**
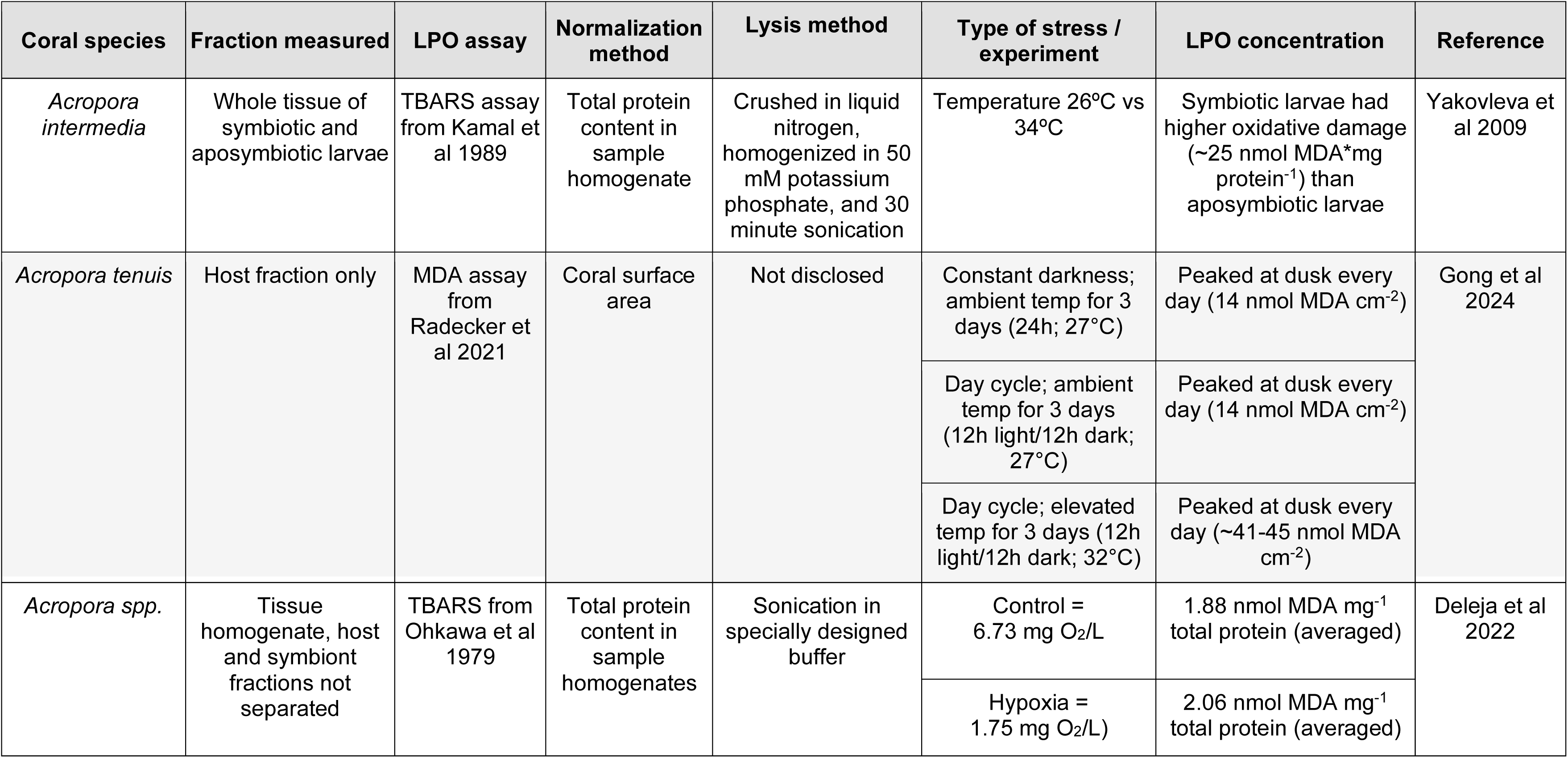
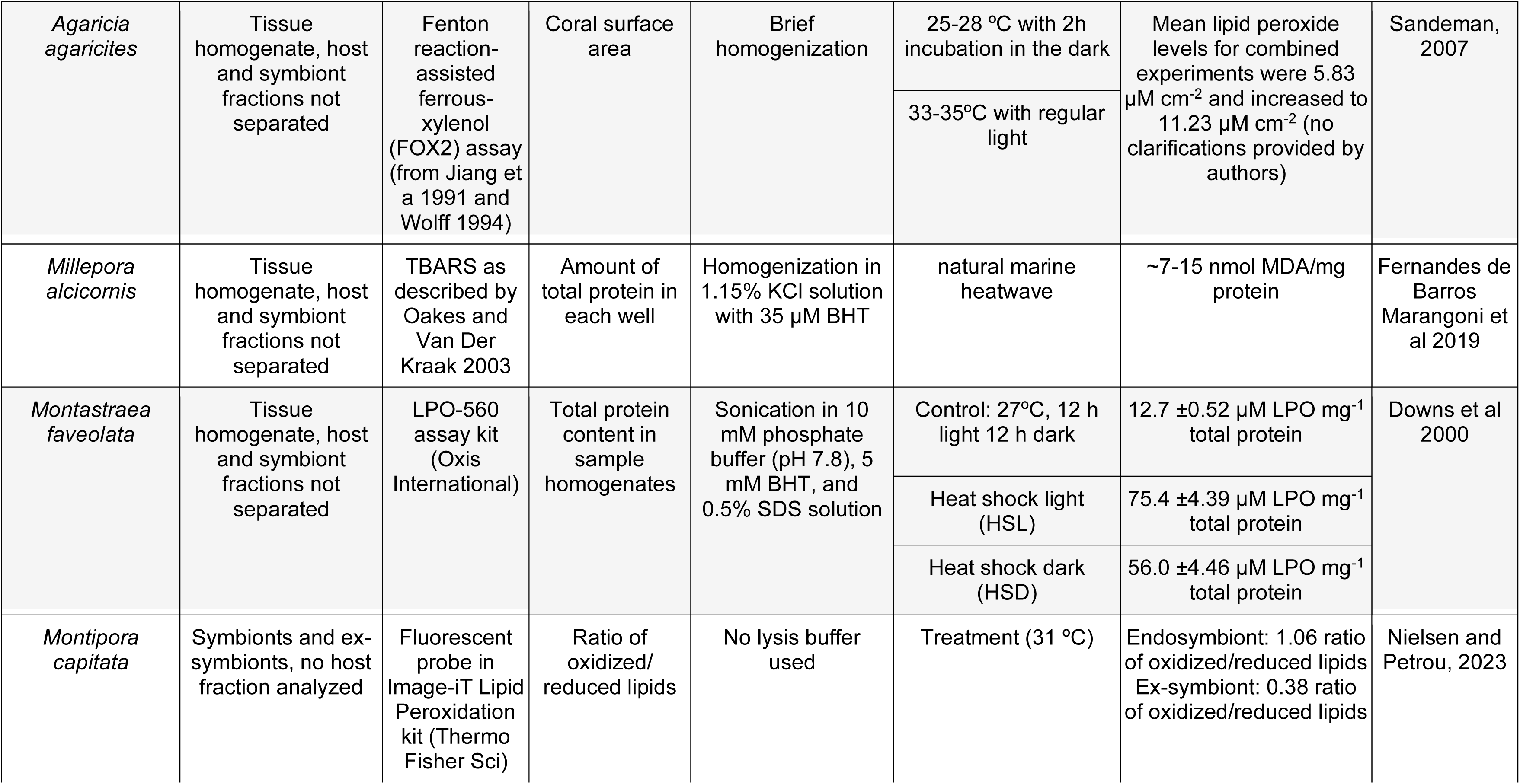
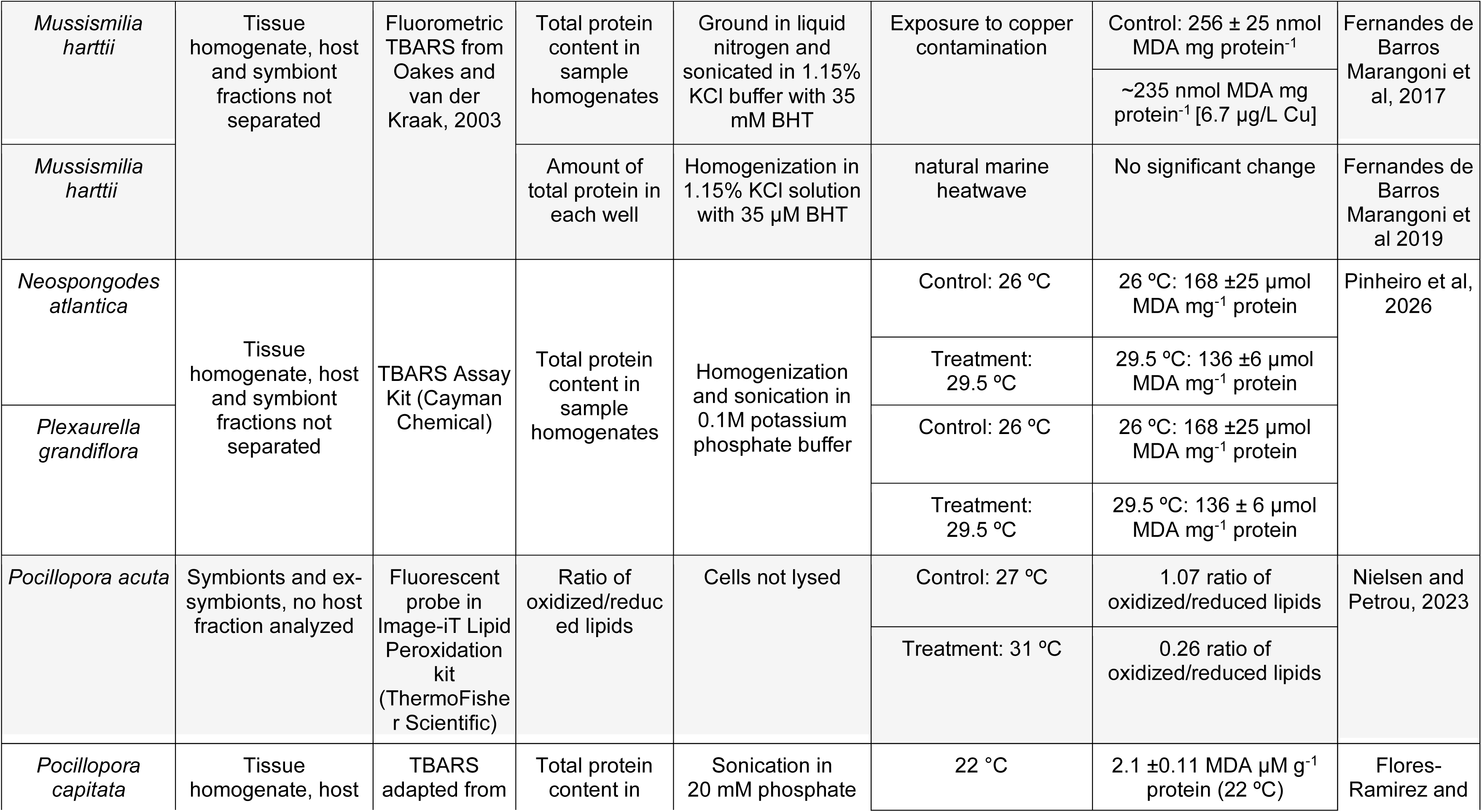
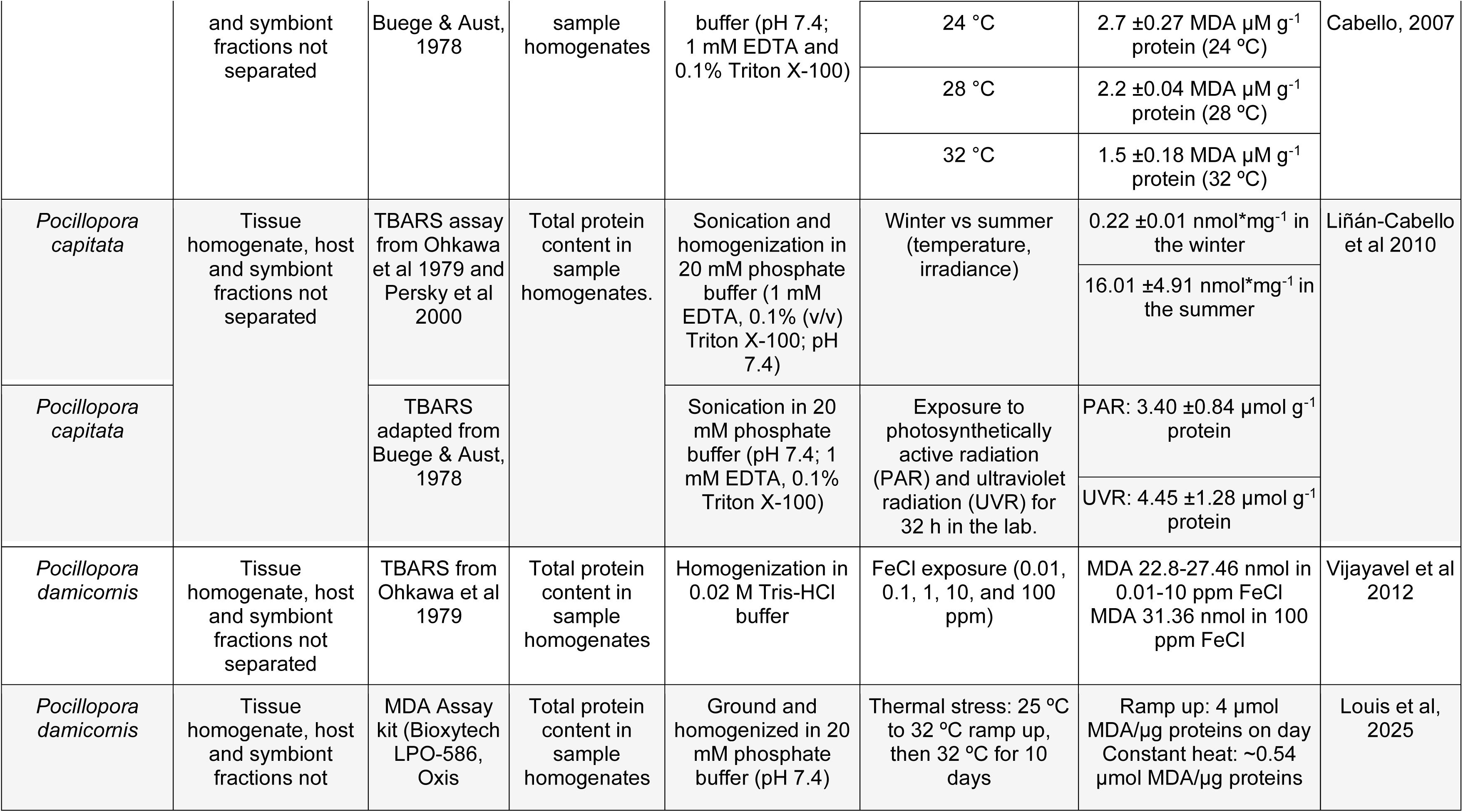
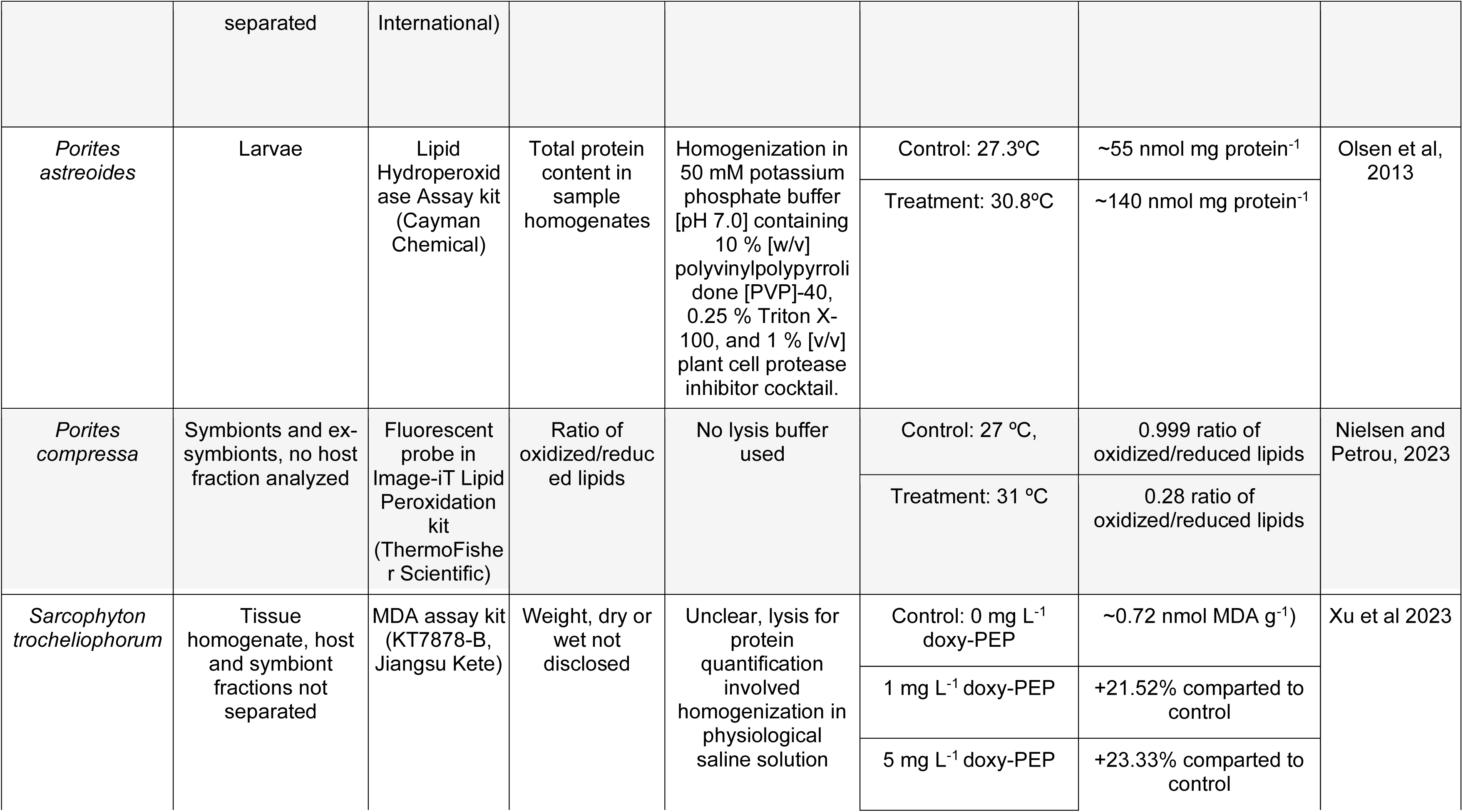
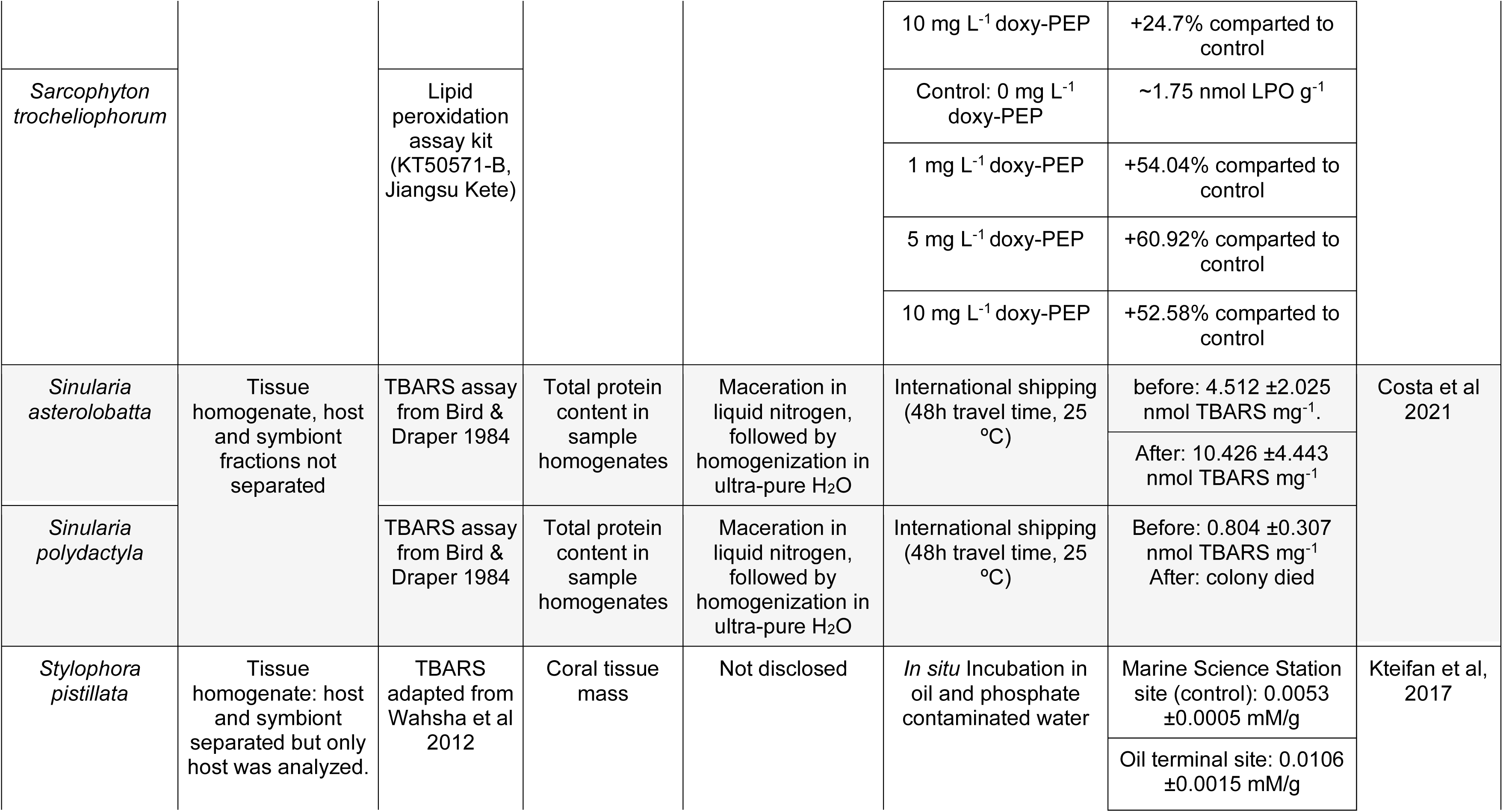
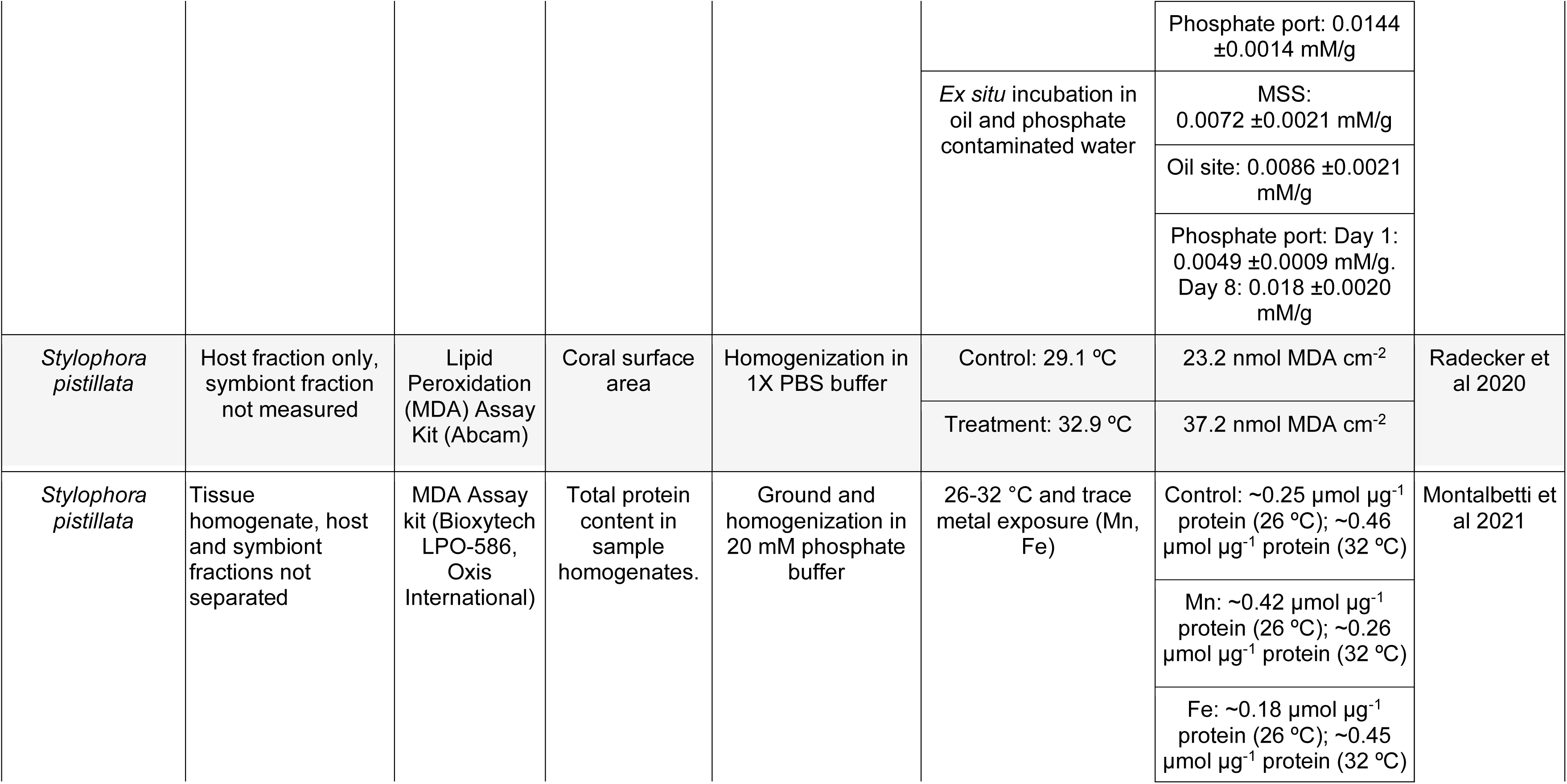
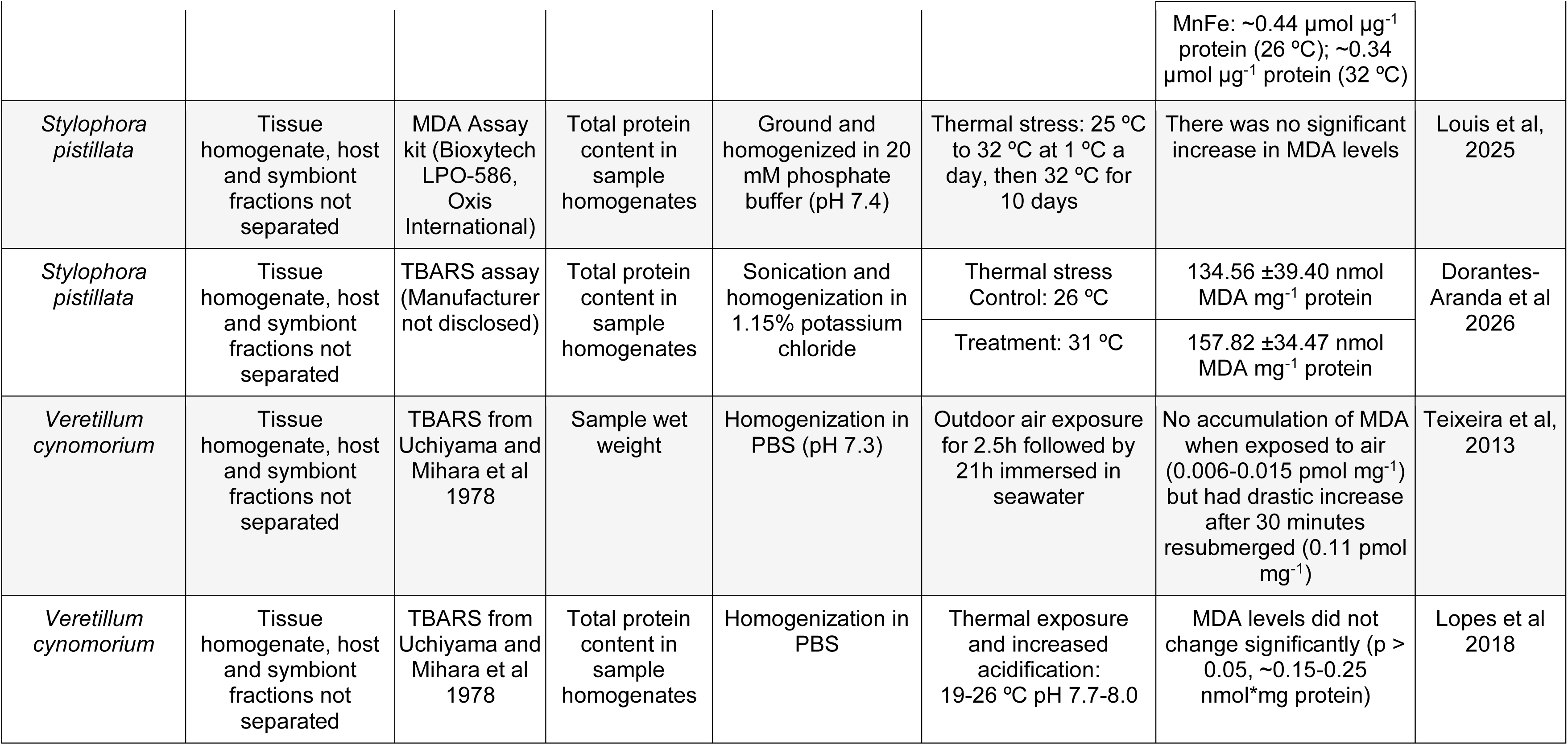
Summary table showing meta analysis of literature reporting on lipid peroxidation in coral bleaching biology.

Key sources of methodological heterogeneity across studies include the coral holobiont fraction analyzed (holobiont versus host and algal symbiont separately), the lysis protocol (including lysis buffer composition), the normalization metric (e.g., whether MDA is normalized to protein concentration, cell count, or surface area), and the chosen assay, as summarized in Table 1. Inconsistency in fraction selection in particular limits the ability to draw universal conclusions about the relationship between LPO and coral bleaching, as unseparated holobiont measurements and fraction-specific measurements are not anticipated to be directly comparable.

A common assay used in coral LPO research is the TBARS assay (Flores-Ramírez & Liñán-Cabello, 2007; Gong et al., 2024; Rädecker et al., 2021; Table 1). It was originally developed for mammalian and food science applications (Aguilar Diaz De Leon & Borges, 2020; Panis et al., 2012, Papastergiadis et al., 2012; Wenjiao et al., 2014), while its utilization in coral tissue requires careful attention to several assay parameters that are rarely acknowledged in the literature. Additionally, MDA quantification alone may provide an incomplete picture of LPO dynamics, because LPO is nonspecific and generates a wide array of products, including lipid hydroperoxides, lipid hydroxides, isoprostanes and neuroprostanes, various aldehydes and ketones, and hydrocholesterols (Niki, 2014). Despite of an incomplete capturing of LPO, the measurement of MDA was found to be a reliable proxy for oxidative stress in mammalian tissue (Aguilar Diaz De Leon & Borges, 2020; Niki, 2014; Panis et al., 2012). Its application to corals is more complex however, owing to the presence of endosymbiotic algal cells. While the coral host cells share the same fundamental animal cell architecture as mammalian cells (nucleus, ER, mitochondria, cytosol; Baumann and Walz, 2001), the algal symbionts introduce a biochemically distinct photosynthetic compartment with its own lipid pool and free radical-generating capacity, a feature absent from mammalian systems and one that fundamentally alters the cellular matrix in which LPO is being measured.

The architecture of algal cells, here dinoflagellates, also presents a practical challenge for lysis. Unlike animal cells, which are bounded by lipid bilayers, dinoflagellates possess a cortical membranous complex called the amphiesma, comprising of alveolar amphiesmal vesicles and an outer plate (Kwok et al, 2023), which confers substantially greater structural rigidity and resistance to lysis. This necessitates lysis optimization, as conditions must be sufficiently stringent to disrupt dinoflagellate cells yet gentle enough to avoid further oxidative damage to host cell membranes. Additionally, animal and algal cells prefer to be in their optimal physiochemical environments, including pH, and temperature, and the coral symbiocyte presents the unusual circumstance of both cell types cohabitating in a seawater-equilibrated milieu (Allemand et al, 2018). These factors introduce potential sources of interference that are absent from mammalian systems and that have received insufficient attention in the coral LPO literature.

Here, we pursue three objectives toward a coral-specific framework for standardized LPO measurement. First, we conduct a systematic review of the existing coral LPO literature to characterize the extent and sources of methodological variation across studies. Second, using our own experimentally derived measurements from a controlled thermal stress experiment on *Montastraea cavernosa*, we apply the TBARS assay separately to host and algal symbiont fractions, generating novel fraction-resolved MDA data that demonstrate both the feasibility and the interpretive necessity of this approach. Third, drawing on both the literature review and our empirical results, we propose minimum reporting guidelines for coral LPO measurements to improve cross-study comparability. Aggregate holobiont LPO measurements conflate two biochemically distinct sources of oxidative stress and cannot be assumed to reflect host- or symbiont-specific dynamics without fraction separation. By combining our empirical findings with a systematic review of existing coral LPO studies, we propose a path toward more standardized and interpretable measurement in this system.

## 2. Materials and Methods

### 2.1 Meta analysis

We conducted a systematic literature review of reported coral LPO measurements using Google Scholar to identify peer-reviewed journal articles. The keywords used in the search were ‘lipid peroxidation in corals’, lipid peroxidation coral’, and ‘lipid peroxidation coral biology’. We included papers published within the past 30 years (1996-2026) that conducted assays to report LPO measurements in corals, extracting the following variables where available: fraction analyzed (holobiont or host and algal symbiont separately), the method used to measure LPO (TBARS assay, Fox2 assay, etc), and the stressor used to induce oxidative stress. In total, we identified 23 papers that together illustrate the extent of methodological heterogeneity and reporting discrepancies associated with using LPO as a proxy for oxidative damage in coral stress research.

To provide a comparison, we also collated LPO data from studies conducted on human tissues (Supp. Info. Table S2).

### 2.2 LPO assay development

#### 2.2.1 Coral sample collection and experimental conditions

The coral samples used in this study were derived from an outdoor thermal stress experiment conducted in Bermuda examining pulsed artificial upwelling (AU) as a potential mitigation strategy for thermal stress. Briefly, 6 colonies of *Montastraea cavernosa* were collected from the Bermuda rim reef (Hog Reef; 32.46°N, 64.83°W; Protected Species License No. 2024061212) in July 2024, were fragmented, and allowed to acclimate for two weeks at 28 °C at the Bermuda Marine Mesocosm Facility of the ASU Bermuda Institute of Ocean Sciences. Fragments of each colony were evenly distributed between three different treatments: ambient (28 °C), heat (30.5 °C), and heat_AU (30.5°C / 27.5°C). Heat_AU simulated upwelling of 24 °C-water for 8 hours each day (8:00 to 16:00) decreasing temperature to 27.5°C during that time.

Samples for LPO analysis were collected on day 22 of the experiment (15 August 2024): one fragment per colony per treatment at four intra-day time points (08:00, 13:00, 17:00 and 20:00), snap-frozen in liquid nitrogen and stored at -80 °C. The tissue was removed from the skeleton by air picking (high-pressure air-gun) in a small amount of chilled 45 μm-filtered seawater. The resulting tissue slurry was subsequently transported in a dry shipper at −120 °C from BIOS to Arizona State University and stored at -80 °C until lipid peroxidation analysis.

#### 2.2.2 Host and algal symbiont fraction separation

Host and algal symbiont fractions were separated through differential centrifugation prior to lysis. Briefly, 500 μL of coral slurry was homogenized on ice for 2 minutes using a handheld homogenizer (Fisherbrand 150 Handheld Homogenizer, Plastic Disposable Generator Probe) with 5 μL of 100X butylated hydroxytoluene (BHT; from RayBio Lipid Peroxidation (MDA) Assay Kit (Catalog #: MA-MDA-2, RayBiotech, Georgia, USA)) to inhibit further LPO during processing. Samples were centrifuged at 1,000 rcf for 5 minutes at 4 °C, pelleting the algal symbiont fraction. The supernatant was retained as the host fraction, and the pellet was resuspended in 500 μL of filtered sterile artificial seawater (FSASW) and centrifuged again under the same conditions to minimize host contamination of the algal symbiont fraction. The supernatant from this wash step was discarded prior to lysis.

### 2.2.3 Lysis and MDA quantification

Both fractions were lysed using a sodium phosphate lysis buffer developed specifically for this study (50 mM sodium phosphate, pH 7.4, 150 mM NaCl, 0.05% w/v TritonX 100). Since LPO-associated compounds are highly labile and LPO itself can be artificially induced by mechanical or chemical perturbation during processing, we prioritized a lysis approach that minimized additional oxidative insult to samples that had already undergone thermal stress and cryogenic preservation. Mechanical methods such as mortar and pestle homogenization were incompatible with our air-picked, fraction-separated samples, and sonication at effective lysis intensities for extended periods of time risked both heat-induced LPO and degradation of MDA adducts. We therefore developed a detergent-based buffer optimized for gentle yet effective lysis of both cell types. Most lysis buffers contain detergents to facilitate the chemical breakdown of the cell membrane, but many common detergents are incompatible with the TBARS assay. For example, given its chemical structure, Tris has the potential to react with MDA and MDA-DNA adducts (Niedernhofer et al., 1997), reducing MDA-TBA adduct yield and leading to an underestimation of MDA. Triton-X100 does not react directly with MDA and has been used in buffers for the MDA assay previously (Flores-Ramirez and Liñan-Cabello, 2007; Liñan-Cabello et al, 2010); however, we reduced its concentration relative to prior formulations, as high Triton-X100 concentrations have been reported to artificially elevate LPO (Parwin et al, 2019). The buffer was prepared at 2X concentration to account for dilution introduced by the host supernatant. For algal symbiont samples, 500 μL of 1X lysis buffer was added with 5 μL of BHT. For the host samples, 250 μL of host supernatant was mixed with 250 μL of 2X lysis buffer and 5 μL BHT. Host samples were rocked gently on a shaker plate in a cold room at 4 °C for 30 minutes. Algal symbiont samples were sonicated (QSonica sonication probe, model CL-18) at 30% amplitude for six 15-second pulses with 5-second intervals. All samples were then centrifuged at 14,000 rcf at 4 °C for 10 minutes and the post-lysis supernatant was used directly in the MDA assay.

MDA concentration was measured using the RayBio Lipid Peroxidation (MDA) Assay Kit (Catalog #: MA-MDA-2, RayBiotech, Georgia, USA) according to the manufacturer’s protocol. Critically, all MDA standards were prepared in 1X lysis buffer for matrix matching, a step necessary to account for the novel lysis buffer composition and minimize matrix effects on absorbance readings. 200 μL of each sample or standard was combined with 200 μL of manufacturer-provided sodium dodecyl sulfate (SDS) solution, incubated for 5 minutes at room temperature (22 °C), then combined with 500 μL of TBA solution and incubated for 60 minutes at 95 °C. Following incubation, samples were cooled on ice for 5 minutes and centrifuged at 1,600 rcf for 10 minutes to pellet any precipitate. Triplicate aliquots of each supernatant were transferred to a clear, flat-bottomed 96-well plate and absorbance was read at 532 nm using a Cytation 5 microplate reader and imager (BioTek Agilent).

Absorbance values were blank corrected, and a linear regression of the standards was performed with a y-intercept set at 0. Individual absorbance readings were converted to MDA concentration in nmol MDA mL^-1^. Host concentration was adjusted for dilution.

#### 2.2.4 Normalization

MDA concentrations were normalized to total protein concentration in nmol MDA mg^-1^ protein to account for differences in biomass across samples, a normalization choice that is itself variable across coral LPO literature and is therefore discussed further in the context of methodological standardization later in the text. Protein concentration was determined using the Bicinchoninic Acid Assay (BCA, Pierce™ BCA Protein Assay Kit, ThermoScientific, 23225) following the manufacturer’s protocol with some adaptions. The BCA reagent was prepared at a 50:1 ratio of the manufacturer’s Reagent A and Reagent B. Bovine Serum Albumin (BSA) standards were prepared from the kit-provided 2,000 μg/mL stock and plated in triplicate (25 μL), along with a triplicate 1X lysis buffer blank for background subtraction. Post-lysis supernatant from each sample was likewise plated in triplicate. Importantly, samples intended for the BCA assay were processed without BHT to prevent potential interference with the colorimetric reaction. 200 μL of BCA working solution was added to each well, mixed gently, and the plate was incubated in the dark at 37 °C for 30 minutes. The plate was then cooled at room temperature for 10 minutes before absorbance was read at 562 nm (Cytation 5, BioTek Agilent).

Protein concentrations were determined following blank subtraction of MilliQ water from the standards and 1X lysis buffer from the samples. Standard replicates were averaged and a linear regression was taken with a set intercept at (0,0), and background-subtracted absorbance readings were set equal to the standard curve. Host absorbance measurements were corrected for dilution. A separate calculation was performed to normalize MDA concentration to coral surface area instead of protein concentration in order to provide additional comparison.

#### 2.2.5 Statistical analysis

Statistical analysis of MDA concentrations was performed with the software R (v. 4.5.2; R Core Team) via linear mixed effects models using the *lmer* function (*lme4* package). MDA concentration normalized to protein concentration was modeled as a function of treatment, time of day, and the interaction between treatment and time of day, with colony as a random effect. Using the *emmeans* package, post hoc pairwise comparisons were performed for treatments within timepoints and timepoints within treatments. Comparisons were controlled with Tukey-adjusted p-values. Significance letters for the summarization of significant differences for treatments within timepoints were determined using the *cld* function in the *emmeans* package (Sidak adjustment). Data were visualized using *ggplot2*. Estimated marginal means and 95% confidence intervals were calculated using *emmeans*.

## 3. Results

### 3.1 Literature review

Our systematic review identified 23 papers reporting LPO measurements in coral under stress research, comprising 30 distinct data entries across different species, stressors, or experimental conditions (Table 1). A large majority of entries (77% of entries, 74% of studies) measured LPO in whole holobiont or undifferentiated tissue homogenate, without separating host and algal symbiont fractions. Of the remaining entries that did separate fractions in some form, three reported host tissue only, three reported symbiont or ex-symbiont fractions only, and just one study (*Agaricia agaricites*; Jiang et al 1991) reported both host and symbiont fractions from the same experiment.

Assay choice was similarly dominated by a single method: a TBARS- or MDA-based colorimetric assay (80% of entries), while the remainder used a Fenton reaction-based FOX2 assay, a fluorescent oxidized/reduced lipid probe, or a lipid hydroperoxide kit. Despite this apparent consistency in assay type, normalization metrics varied considerably: over half of the studies normalized LPO to total protein content, three studies to coral surface area, three to a ratio of oxidized to reduced lipids, and the remaining four studies to sample wet weight, dry weight, or whole-tissue mass. Lysis protocols showed comparable heterogeneity, ranging from brief manual homogenization to multi-step sonication in buffers differing in composition, antioxidant additives (e.g., BHT), and pH, with several studies not disclosing lysis conditions at all (Table 1).

Together, these patterns indicate that the coral LPO literature is dominated by aggregate holobiont measurements, relies overwhelmingly on a narrow set of TBARS/MDA-based assays, but lacks consensus on how LPO should be normalized or how tissue should be prepared prior to analysis. These are the three independent axes of methodological variation that limit direct comparison across studies.

### 3.2 Heat stress experiment

Under ambient conditions, mean algal symbiont MDA levels were 40.3% lower than mean host MDA levels (52.7 ±12.9 vs. 88.3 ±7.64 nmol MDA mg⁻¹ protein, respectively; Figure 1). Both fractions exhibited non-significant diel variation in MDA, though the pattern was more pronounced in the algal symbiont fraction. Host MDA increased by 12.1% from 08:00 to 20:00 in the ambient treatment, and algal symbiont MDA increased by 49.4% from 08:00 to 13:00 (42.7 ±14.9 to 63.9 ±14.4 nmol MDA mg⁻¹ protein) and subsequently decreased by 7.5% at 17:00 (59.1 ±14.6 nmol MDA mg⁻¹ protein) and by a further 23.8% at 20:00 (45.0 ±14.4 nmol MDA mg⁻¹ protein).

**Fig. 1.**
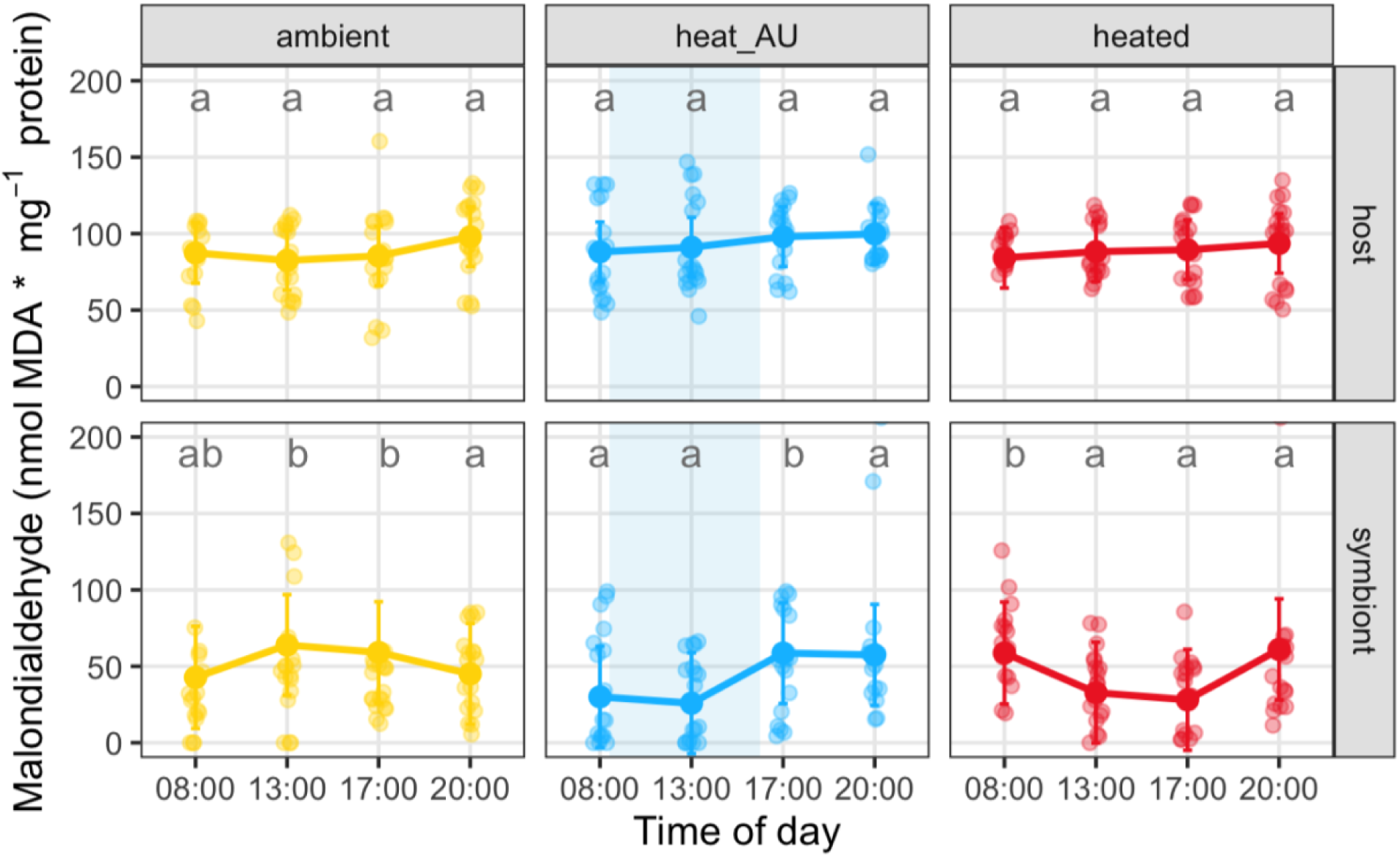
MDA concentration in *M. cavernosa* (nmol MDA mg^-1^ protein) after 22 days of exposure to ambient, heat_AU (artificial upwelling for 8h starting at 8:00 am), and heated treatments in the host and algal symbiont fractions. Opaque circles represent the mean MDA concentration, and partially transparent circles represent individual data points and replicates. Error bars depict standard error. The light blue shaded area indicates the cold water pumping period in the heat_AU treatment. Letters indicate significant differences in pairwise, post hoc comparisons for treatments within timepoints

Under heat-stress conditions, host MDA (88.9 ±7.60 nmol MDA mg⁻¹ protein) did not differ significantly from the ambient condition. In contrast, algal symbiont MDA was significantly lower than ambient at 13:00 (32.8 ±14.5 nmol MDA mg⁻¹ protein; p = 0.0034) and 17:00 (28.1 ±14.5 nmol MDA mg⁻¹ protein; p = 0.0050). The diel trajectory in the heat-stress algal symbiont fraction began at 58.7 ±14.9 nmol MDA mg⁻¹ protein at 08:00, declined by 44.2% at 13:00, and declined further by 14.3% at 17:00 (cumulative decline from 08:00 to 17:00: −52.1%, p = 0.040). A significant rebound of +117% was then observed at 20:00 (61.1 ±14.6 nmol MDA mg⁻¹ protein; p = 0.018).

Mean host MDA in the heat_AU treatment (94.2 ±7.60 nmol MDA mg⁻¹ protein) was slightly, though non-significantly, elevated relative to both ambient and heat-stress treatments. Heat_AU algal symbiont MDA was significantly lower than ambient at 13:00 (25.9 ±14.5 nmol MDA mg⁻¹ protein; p = 0.0002) and significantly lower than heat-stressed algal symbionts at 08:00 (30.0 ±14.5 nmol MDA mg⁻¹ protein; p = 0.0131), but significantly higher than the heat-stress treatment at 17:00 (58.7 ±14.5 nmol MDA mg⁻¹ protein; p = 0.0043). Over the course of the day, heat_AU agal symbiont MDA increased by 13.9% from 08:00 to 13:00, then by 127% from 13:00 to 17:00 (p = 0.017), and remained approximately stable at 20:00 (57.4 ±14.6 nmol MDA mg⁻¹ protein; −2.1% from 17:00; p = 0.025 for 13:00 vs. 20:00).

MDA measurements normalized to coral surface area (Supp. Info Fig. S3) show similar diurnal trends to MDA measurements normalized to protein concentration. Host samples showed no significant differences between treatments in nmol MDA cm^-2^ concentration for 08:00, 13:00, and 20:00, but the heat_AU treatment showed a significantly greater mean MDA compared to the ambient treatment at 17:00 (103.4 ±9.47 nmol MDA cm^-2^; p = 0.0077). Algal symbiont MDA normalized to coral surface area showed steady mean values within the range of 5-11 nmol MDA cm^-2^. The only difference in inter-treatment response compared to MDA normalized to protein concentration was found at 08:00 where mean MDA in the heated treatment showed a statistically significant variation compared to the ambient treatment (9.23 ±2.07 nmol MDA cm^-2^, +61%; p = 0.0026), while heat_AU was not significantly different from either of them.

Individual colony variation in MDA concentration is presented in Figure 2, with each colony showing distinct host and agal symbiont responses to treatments.

**Fig. 2.**
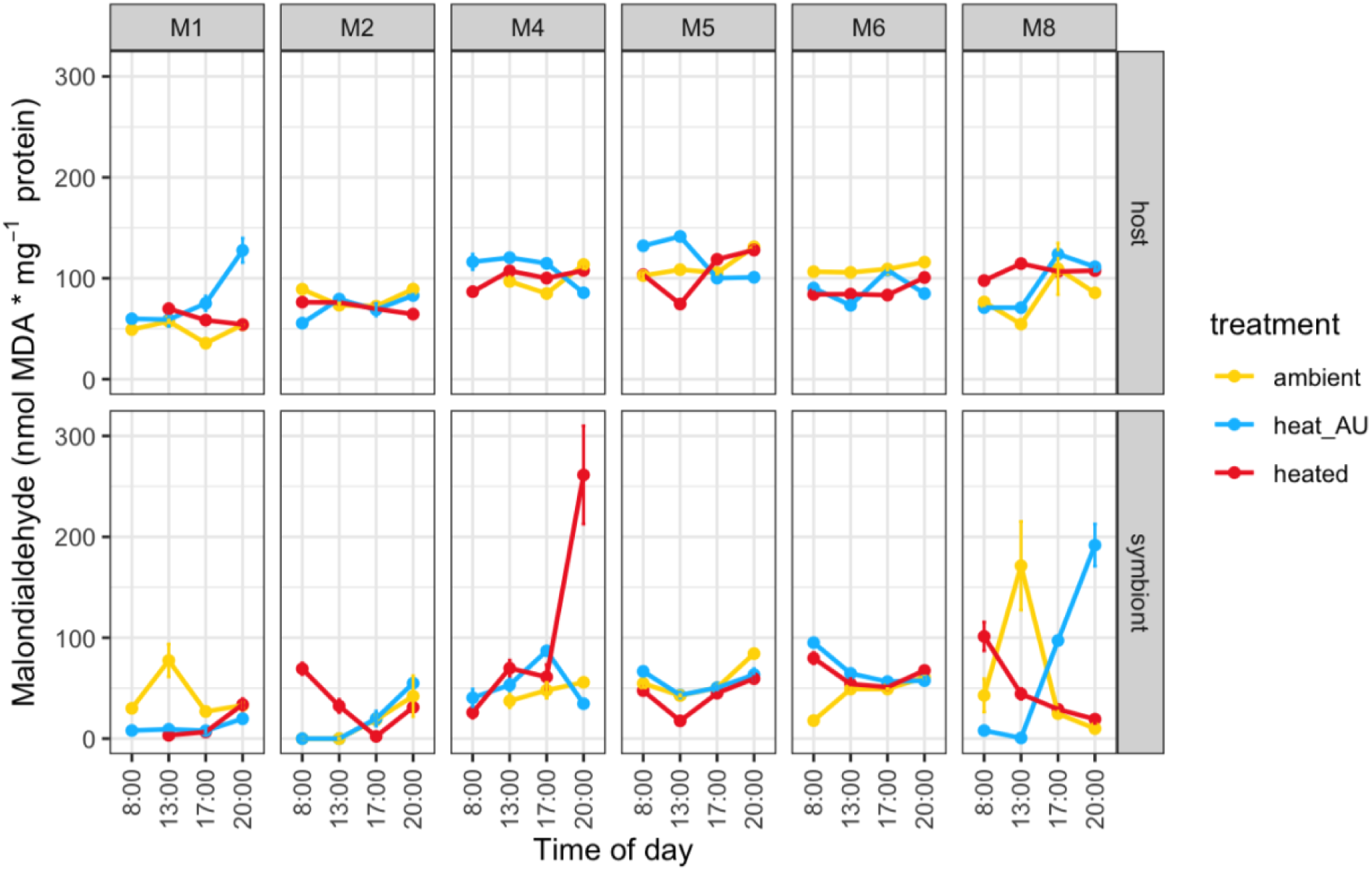
Mean MDA concentrations in the six different colonies of *M. cavernosa* (nmol MDA mg^-1^ protein) after exposure to the ambient (yellow circles and lines), heat_AU (blue circles and lines), and heated (red circles and lines) treatments in the host (top panels) and algal symbiont fractions (lower panels)

An additional factor explored, although not originally intended, is the consistency of the standard curve. Standard curve slopes varied substantially across assays (Figure 3; Supp. Info. Table S4). The two assays performed on the days the standard stock was first opened yielded nearly identical slopes (3.34 and 3.25 mM^-1^ MDA), indicating good inter-day reproducibility for freshly opened standards. Subsequent assays using previously opened stock produced progressively divergent slopes (4.66, 16.59, and 10.91 mM⁻¹ MDA for Assays 3, 4, and 5, respectively), consistent with degradation of the standard stock after opening. To ensure analytical consistency, all sample MDA concentrations were calculated using the standard curves from Assays 1 and 2.

**Fig. 3.**
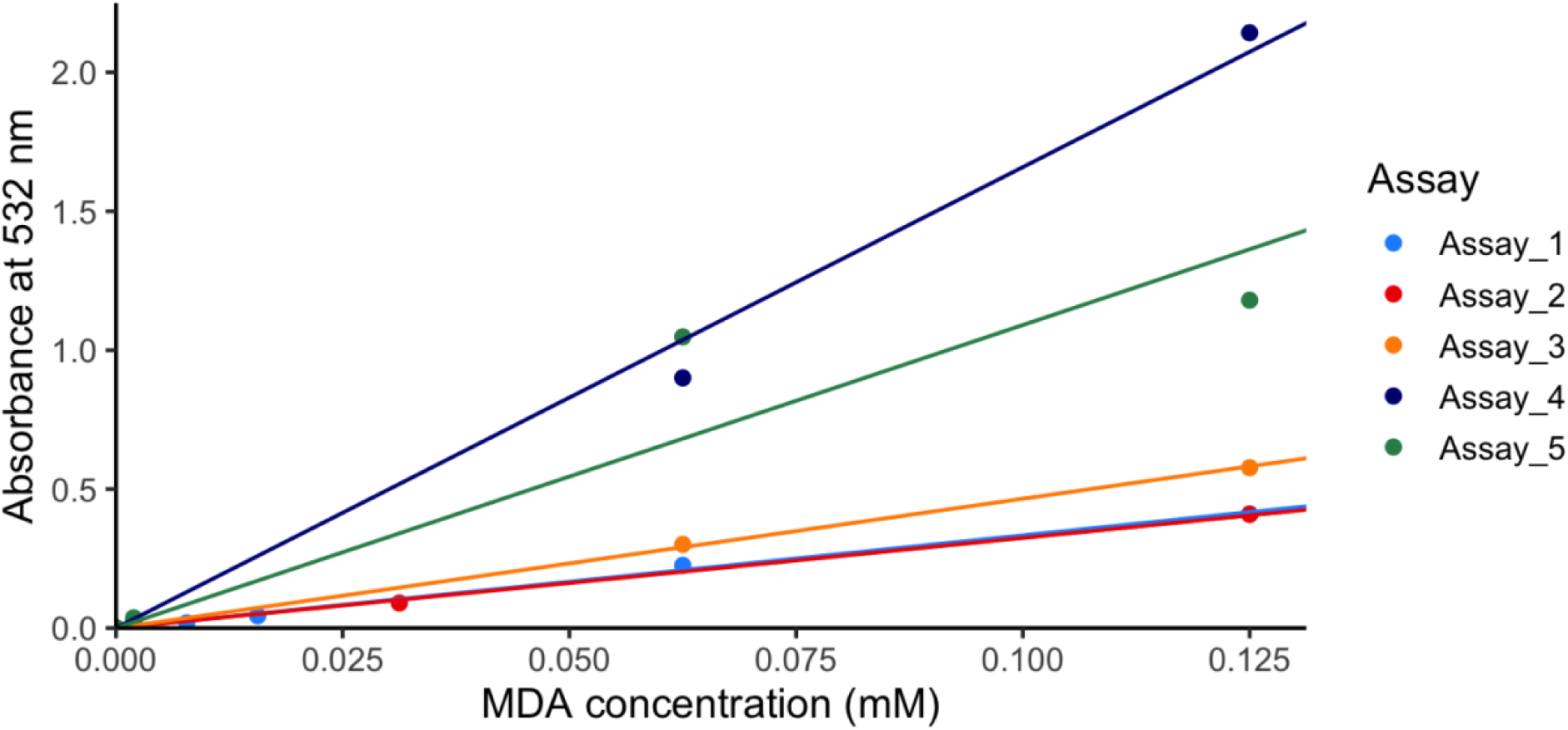
MDA standard curve from different assay runs performed on 5 different days. Absorbance was blank subtracted prior to plotting. Individual data points represent the specific absorbance readings of each standard, and solid lines represent the regression lines obtained from each assay run. The standard concentrations used for each assay are as follows: Assay 1: 0 mM, 0.00195 mM, 0.00781 mM, 0.01563 mM, 0.0625 mM, 0.125 mM; Assay 2: 0 mM, 0.00195 mM, 0.03125 mM, 0.125 mM; Assay 3: 0 mM, 0.00195 mM, 0.0625 mM, 0.125 mM; Assay 4: 0 mM, 0.00195 mM, 0.0625 mM, 0.125 mM; Assay 5: 0 mM, 0.00195 mM, 0.0625 mM, 0.125 mM

## 4. Discussion

Our experiment demonstrates that host and algal symbiont LPO dynamics are decoupled under thermal stress, and that the choice of normalization metric can itself shift the pattern observed in a given fraction. Below, we first discuss these experimental findings and their implications for interpreting coral oxidative stress, before broadening the discussion to our systematic review of the coral LPO literature, where we argue that fraction separation and normalization are two of several methodological choices currently limiting comparability across studies.

### 4.1. Thermal stress experiment MDA data

The substantially higher baseline LPO in the host fraction relative to the agal symbiont fraction under ambient conditions (88.3 vs. 52.7 nmol MDA mg⁻¹ protein; Figure 1) underscores that the two compartments operate under distinct basal oxidative regimes. This difference, and the fact that the coral host constitutes a larger fraction of the biomass than the algal symbionts, demonstrates that measuring LPO at the holobiont level conflates biochemically distinct signals in the two compartments, as it largely represents the dynamics in the host. The greater colony-to-colony variability in algal symbiont LPO, reflected in consistently larger standard errors, may suggest that algal symbionts are more sensitive to individual colony characteristics (e.g., algal symbiont clade identity, colony microenvironment, symbiont density) than the host tissue under basal conditions.

The absence of significant differences in host LPO between heated and ambient treatments may indicate that the 2.5 °C temperature elevation applied here (28 to 30.5 °C) was either insufficient to elicit a detectable host LPO response over the experimental timeline, or that the corals had acclimatized to a stable, higher temperature over the prior 21 days. In contrast, the agal symbiont fraction exhibited significant treatment differences and a pronounced diel pattern. The lower algal symbiont LPO in the heated treatment at 13:00 and 17:00 relative to ambient, contrary to the expectation of elevated oxidative stress under increased temperature and irradiance (Lesser, 1996), is surprising. The overall diel trajectory of higher LPO at 08:00 and 20:00 and lower LPO during peak irradiance hours suggests that oxidative damage may accumulate preferentially during nocturnal respiration rather than diurnal photosynthesis, or alternatively that daytime antioxidant upregulation temporarily suppresses LPO. Analogous diel patterns have been reported in thermally stressed plants, in which nocturnal carbon metabolism and mitochondrial free radical production have been implicated (Peraudeau et al., 2015; Impa et al., 2018; Abbas et al., 2024); whether a similar mechanism operates in algal symbionts or whether antioxidant production is simply upregulated warrants further investigation.

The heat_AU treatment presents a more complex picture. Algal symbiont LPO was significantly suppressed relative to ambient at 13:00, during simulated upwelling (08:00–16:00), consistent with the thermal mitigation hypothesis. However, a sharp rebound to levels comparable to heat-stressed controls was observed at 17:00 and persisted to 20:00, after the cold-water pulse had ceased. This rebound may reflect a compensatory oxidative response following the end of thermal buffering, or a lag in free radical scavenging as algal symbiont metabolism adjusts to re-warming.

While diel LPO dynamics in thermally stressed corals have rarely been examined, our ambient and heated results are broadly consistent with those of Downs et al. (2000), who reported elevated LPO in *Montastraea faveolata* under heat shock (33 °C for 12 h) relative to controls maintained at 27 °C, with greater LPO when heat shock occurred during the light period than the dark. The light-dependent amplification of LPO observed by Downs et al. (2000) aligns with our finding of a symbiont-driven oxidative response, given that photosynthetic free radical generation is contingent on illumination. Direct quantitative comparison is not possible, however, as Downs et al. (2000) did not report sampling time points, did not separate host and algal symbiont fractions, and used a different normalization approach, precluding attribution of their signal to either compartment. The temporal dynamics in our experimental treatments would not have been apparent from a single-time-point sampling design, highlighting the value of a finer sampling resolution in coral LPO studies to capture potential signs of thermal acclimation or light intensity effects.

More broadly, the absolute LPO values reported here are difficult to compare to the wider coral literature given pervasive inconsistencies in fraction analyzed, assay choice, normalization metric, and sampling time, the latter of which is rarely reported. Assay differences are particularly consequential: HPLC-based MDA quantification has been shown to be more accurate than colorimetric and fluorometric methods in animal systems (Moselhy et al., 2013), meaning absolute values across methods are not directly equivalent. Nonetheless, intra-study trends remain interpretable, and it is the level of trend rather than absolute values that cross-study synthesis is most tractable.

Our findings highlight how holobiont-level measurements can obscure biologically meaningful signals. Dorantes-Aranda et al. (2026) reported elevated holobiont LPO in heat-stressed (31°C for two weeks after a three-week acclimation at 26 °C) *Stylophora pistillata* relative to controls (26 °C), a temperature range comparable to our experimental conditions. However, without host-symbiont fraction separation, the cellular origin of that signal cannot be determined. The discrepancy between our findings and Dorantes-Aranda et al. (2026) is more likely attributable to the larger thermal differential in their study (∼5°C vs. ∼2.5°C here) and species-specific differences between *S. pistillata* and *M. cavernosa*, rather than to a symbiont-driven holobiont signal, since host LPO in our data exceeded symbiont LPO under ambient conditions, and the host constitutes the larger share of holobiont biomass. Where *S. pistillata* host and algal symbiont fractions respond differently from those of *M. cavernosa* under equivalent thermal stress remains an open question, but one that can only be addressed through fraction-resolved measurements of the kind presented here.

Colony-level variation (Figure 2) reinforces that oxidative stress responses in *M. cavernosa* are not uniform across genotypes. Distinct LPO trajectories among colonies in both fractions suggest that genetic background, prior thermal history, or algal symbiont clade may modulate LPO dynamics independently of treatment. These findings underscore the importance of sampling from multiple colonies and treating colonies as a random effect in statistical models.

### 4.2. Standard curve instability and implications for MDA quantification

Despite maintaining consistent experimental procedures across assays, we observed a marked increase in standard curve slope with successive openings of the MDA standard stock solution (Figure 3; Supp. Info. Table S4). Assays 1 and 2, performed on the days the stock was first opened, yielded closely matched slopes (3.34 and 3.25 mM⁻¹ MDA), while 3–5 produced substantially steeper slopes, peaking at 16.59 mM⁻¹ MDA in Assay 4. This pattern is most consistent with progressive degradation of the MDA standard after opening, potentially through oxidation, polymerization, or solvent evaporation, resulting in elevated background absorbance and apparent hypersensitivity of the standard curve. This hypersensitivity could lead to an underestimation of LPO levels in certain samples tested with subsequent standard openings. The non-monotonic decrease from Assay 4 to 5 (16.59 to 10.91 mM⁻¹) may reflect partial re-equilibration or micro pipetting variation but does not invalidate the overall pattern. No indication of standard degradation or specific preservation methods were mentioned in the manufacturer’s instruction manual. While we used only one commercial kit, the MDA–TBA chemistry is common across TBARS-based kits, and similar degradation behavior is plausible for other commercial standards.

To ensure analytical consistency in the present dataset, all sample MDA concentrations were calculated against the standard curves from Assays 1 and 2. This is acknowledged as a limitation: samples measured in Assays 3–5 were back-calculated using curves that may not perfectly reflect those assay-day conditions.

### 4.3 Sources of methodological variation in coral LPO research

The systematic literature review (Table 1) reveals that methodological heterogeneity is pervasive in coral LPO research and is a primary driver of cross-study variation. Four key sources of variation are identifiable: (1) the holobiont fraction analyzed; (2) lysis method and buffer composition; (3) choice of LPO assay; and (4) normalization metric.

Regarding the separation of host and algal symbiont fractions, the majority of studies reviewed measured LPO in bulk tissue homogenates (Table 1), conflating host and algal symbiont signals. As our empirical data demonstrate, the two compartments exhibit divergent basal LPO levels and distinct responses to thermal stress; holobiont-level measurements therefore cannot attribute changes in MDA to a specific partner.

Regarding matrix matching, failure to match the background composition of standards and samples introduces systematic bias into absorbance-based assays. Algal symbiont chlorophyll absorbs near the 532 nm MDA–TBA chromophore peak and can confound readings if not adequately removed during post-lysis centrifugation (Morales and Munñe-Bosch, 2019). Preparing standards in the same buffer matrix as samples, as performed here, is a straightforward measure to minimize this error.

A recognized limitation of the TBARS assay is that TBA reacts with aldehydes beyond MDA, including both LPO-derived and non-LPO compounds whose adducts absorb at overlapping wavelengths, potentially leading to overestimation of MDA (Kosugi et al., 1987; Liu et al 1997; Meras et al 2020). This non-specificity is an inherent constraint of colorimetric TBARS methodology and should be considered when interpreting absolute MDA values, including those reported here.

Normalization heterogeneity further complicates cross-study comparison. MDA values from literature (Table 1) are reported in units spanning nmol mg⁻¹ protein, nmol cm⁻², µmol g⁻¹ wet weight, and ratios of oxidized: reduced lipids, units that are not interconvertible without additional information. We recommend that future studies report LPO normalized to both protein concentration and surface area (or cell count for isolated algal symbionts) to provide baseline units, facilitating comparison across methodological frameworks, while acknowledging that neither metric is without limitation.

Protein concentration normalizes MDA to biomass and scales with both cell size and cell number (Lundberg et al, 2008), making it a more reliable proxy for the amount of biological material being assayed than surface area alone. However, because protein content is also sensitive to cellular metabolic state (Krüger et al, 2012; Petrou et al., 2018), thermal stress itself may alter protein concentration, potentially confounding normalization. Surface area normalization is similarly imperfect: it does not account for tissue thickness or density, and is not directly comparable across coral growth forms, as perforate and imperforate corals differ substantially in the amount of tissue per unit skeletal surface area. Since both metrics are susceptible to stress-induced variation and neither fully captures the biological quantity of interest, reporting both independently is preferable to relying on either alone, and any divergence between the two should be acknowledged and interpreted rather than ignored.

### 4.4 Proposed LPO measurement and reporting framework

To address the sources of variation identified above, we propose minimum reporting and protocol guidelines for LPO studies in corals (Table 2). Central to these recommendations is the separation of host and algal symbiont fractions prior to analysis, which we argue is necessary for interpretable LPO measurement regardless of the biological question under investigation, for the reasons outlined above. We further recommend that researchers report: the holobiont fraction analyzed and the fraction separation method employed; lysis method and, where applicable, buffer composition (concentration, pH) and detergent concentration; the specific LPO assay used (including manufacturer and catalogue number); and experimental design details including the number of colonies, replicates, and sampling time points. Reporting of sampling time point is particularly important given the diel dynamics documented here. Where commercial kits are used, we additionally recommend recording the lot number, the number of times standard stock solutions have been opened, and the elapsed time since first opening, as standard instability can introduce substantial within-study variation that is rarely acknowledged. Additionally, we recommend that researchers prepare fresh MDA standard stocks from solid MDA–TBA adduct or TEP for each assay where possible. Adoption of these guidelines would substantially improve the comparability of coral LPO data across studies and laboratories.

**Table 2.** Suggested minimum reporting guidelines for studying LPO in corals.

| Category | Reporting element | Description |
| --- | --- | --- |
| Organism | Species identification | Taxonomic identification of coral species as well as known symbionts. |
|  | Collection data | Location (coordinates), date, time of day, depth, permits (if applicable) |
| | Environmental conditions | Temperature (°C), salinity (PSU), irradiance (PAR in $\mu\text{E m}^{-2} \text{s}^{-1}$), pH |
| Experimental design | Controls | Which controls were used in the experiment, and corresponding LPO values |
|  | Stressors | Experimental independent variables and their administration |
|  | Assay replication | The number of replicates for each sample in the assays performed |
|  | Colonies used | The number of colonies used from a particular species |
|  | Experimental replicates | The number of replicates in the experimental design, if different from number of colonies |
| Fraction separation | Fraction analyzed | Whether results are reported for the host only, symbiont only, or the unseparated holobiont |
|  | Separation method | How cells were separated, if they were separated |
| Cell lysis | Lysis buffer chemistry | Composition of buffer, concentration, pH, whether this was different between the fractions. If no lysis buffer is used, confirmation of that should be included |
|  | Lysis method | Any mechanical methods used to lyse the cells (e.g. sonication, homogenization), and whether this differed between fractions |
|  | Potential interference | Report any potential reactions between the lysis buffer reagents and experimental reagents which may affect reads. Also mention any other ways in which the lysis method may contribute to skewed readings, and how this was minimized |
| LPO measurement | Assay used | Name of assay, name of manufacturer if purchased, or full description of assay methodology if not purchased from a manufacturer |
|  | Assay chemistry | How the assay works (colorimetric, etc) and how the particular method used may affect this |
|  | Matrix matching | Report attempts made to matrix match the standards and blanks to sample composition |
|  | LPO standards | Date opened, date prepared, elapsed time since opening, standard curve slopes |
| Data analysis | Normalization | LPO measurements normalized to two minimum methods: protein concentration, weight and/or surface area, independently |
|  | Reporting units | Report values in nmol (TBARS or MDA) cm<sup>-2</sup> or nmol (TBARS or MDA) mg<sup>-1</sup> protein |

## 5. Conclusion

This work originated from our empirical LPO measurements, during which we encountered the full range of methodological challenges documented here. These challenges and the difficulty of contextualizing our results against a literature rife with methodological inconsistency, revealed a pressing need for standardization that we address by pairing our empirical findings with systematic review of existing coral LPO studies to propose a path forward. Our systematic review demonstrates that the coral LPO literature is characterized by pervasive methodological heterogeneity in fraction separation, lysis, assay choice, and normalization, heterogeneity that makes it difficult to determine whether differences in reported MDA values reflect true biological variation or experimental artifacts. Our empirical results demonstrate that host and algal symbiont LPO are decoupled: host MDA was stable across treatments, while algal symbiont MDA exhibited significant treatment-dependent and diel dynamics that were only detectable through fraction-specific measurement. We further document an underreported practical concern: commercial MDA standard stocks appear to degrade after opening, with progressive increases in standard curve slope that can substantially distort concentration estimates. Future studies should separately quantify host and algal symbiont LPO, adopt matrix-matched standards, and report the methodological details specified in Table 2. More broadly, expanding LPO studies to encompass multiple species, agal symbiont clades, thermal histories, and diel time points will be essential for understanding the mechanistic role of oxidative membrane damage in coral bleaching and for evaluating the effectiveness of reef conservation strategies.

## Supporting information

all supp mat combined

## Acknowledgments

We would like to thank the technicians and undergraduate students involved in the maintenance of aquaria and organisms.

## Author Contributions

S.W. Mastorakos: Data curation, formal analysis, investigation, methodology, software, visualization, writing - original draft, writing - review and editing.

A.J. Kruger: Data curation, investigation, methodology, project administration, writing - original draft, writing - review and editing.

C. Carbonne: Data curation, formal analysis, methodology, resources, software, validation, writing – review and editing

Y. Sawall: Funding acquisition, resources, validation, writing - review and editing.

L.M. Roger: Conceptualization, funding acquisition, methodology, project administration, resources, validation, visualization, writing - original draft, writing - review and editing.

## Data availability

The detailed protocols for sample fraction separation and the adapted MDA assay are available on Protocol.io.

(Sample fraction separation: https://www.protocols.io/private/C61AA4CD643611F1A7480A58A9FEAC02)

(MDA Assay: https://www.protocols.io/private/87CBCD5A637D11F1A7480A58A9FEAC02)

