## Supplementary material for "Improving coral oxidative stress assessments through compartment-specific lipid peroxidation measurements and increased methodological standardization": all supp mat combined

### Holobiont lipid peroxidation measurements obscure compartment-specific oxidative stress in corals

ORCID ID: 0000-0003-2274-8311

**List of supporting information:**

**S1.** Lipid peroxidation mechanism schematic.

**S2.** Table showing meta-analysis of human LPO levels.

**S3.** Summary of MDA content in coral cells normalized to surface area.

**S4.** Table showing contradicting MDA standard curves.

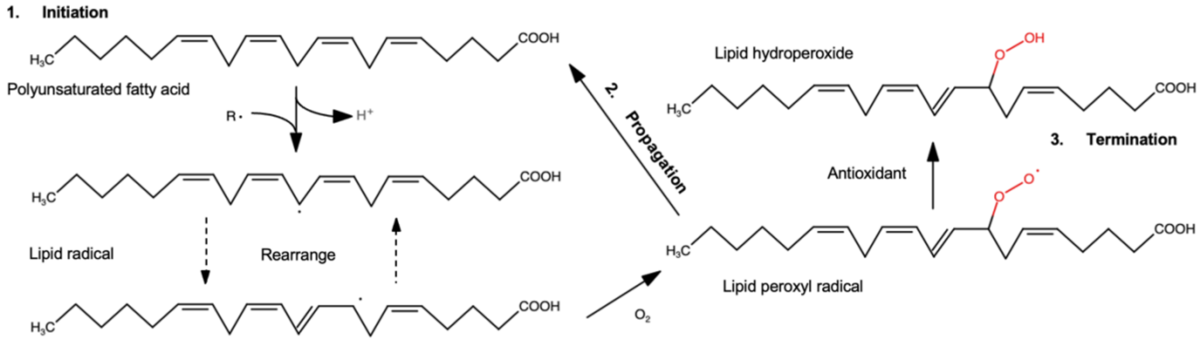


**Fig. S1** Lipid peroxidation mechanism schematic beginning with initiation, moving to propagation, and ending with termination

**Table S2.** Summary table showing literature review on articles that analyzed LPO in human tissue.

| *Species* | Tissue type | Assay used | LPO | Type of stress | Normalization method | Reference |
|---|---|---|---|---|---|---|
| *Homo sapiens* | Whole blood | TBARS assay (spectrophotometric) as in Oliveira et al 2000 | ED: (155.8 ±30.46 nmols/L) AD: (166.7 ±19.86 nmols/L) controls (101.6 ±7.06 nM), | Breast cancer disease progression in early disease (ED) and advanced disease (AD) patients. | Normalized to volume of blood sample. | Oliveira et al 2000; Panis et al 2012 |
| *Homo sapiens* | Colorectal tissue and colon musosae (control) | MDA assay as in Ivanovic et al 1999 | Stage II: 1.7 ±0.39 nmol/g tissue MDA Stage III: 2.25 ±0.43 nmol/g tissue MDA Stage IV: 4.04 ±0.47 nmol/g tissue MDA Control: 1.39 ±0.15 MDA | Colorectal cancer progression across stages II-IV. | Normalized to tissue sample weight. | Ivanovic et al 1999; Skrzydlewska et al 2005 |
| *Homo sapiens* | Colorectal tissue and colon musosae (control) | 4-HNE assay as in De Leenheer et al 1979 | Stage II: 0.37 ±0.007 nmol/g tissue 4-HNE Stage III: 0.45 ±0.09 nmol/g tissue 4-HNE. Stage IV: 0.52 ±0.11 4-HNE Control:0.29 ±0.03 nmol/g tissue 4-HNE | Colorectal cancer progression across stages II-IV. | Normalized to tissue sample weight. | De Leenheer et al, 1979; Skrzydlewska et al 2005 |

| | | | | | | |
|---|---|---|---|---|---|---|
| *Homo sapiens* | Blood serum | TBARS assay as in Yagi 1978 | Breast cancer: 133.24 ±12.94 nmol TBARS / 100 mg protein<br>Control: 126.58 ±11.62 nmol TBARS / 100 mg protein | Breast cancer disease compared to healthy patients | TBARS normalized to 100 mg protein | Yagi 1978; Rajneesh et al, 2008 |
| *Homo sapiens* | Blood serum | Conjugated diene (CD) measurements as in Rao and Recknagel 1968 | Breast cancer: 100 mg protein<br>0.61 ±0.05 µmol CD / haemoglobin<br>Control: 0.46 ±0.04 µmol CD / haemoglobin | Breast cancer disease compared to healthy patients | CD normalized to haemoglobin | Rao and Recknagel 1968; Rajneesh et al, 2008 |
| *Homo sapiens* | Blood serum | Lipid hydroperoxide measurement as in Jiang et al 1992 | Breast cancer: 0.59 ±0.09 nmol LOOH / 100 mg protein<br>Control: 0.44 ±0.08 nmol LOOH / 100 mg protein | Breast cancer disease compared to healthy patients | LOOH normalized to 100 mg protein | Jiang et al 1992; Rajneesh et al, 2008 |

| | | | | | | |
|---|---|---|---|---|---|---|
| *Homo sapiens* | Tumor tissue and lung parenchyma | TBARS assay as in Yagi 1978 | Adenocarcinoma: 3.28 ±1.83 nmol TBARS $L^{-1}$ Small cell lung cancer: 2.94 ±1.91 nmol TBARS $L^{-1}$ Squamous cell lung cancer: 2.23 ±1.77 nmol TBARS $L^{-1}$ ( Large cell lung cancer: 1.97 ±1.06 nmol TBARS $L^{-1}$ | Adenocarcenoma, small cell lung cancer, squamous cell lung cancer, and large cell lung cancer | Normalized to sample volume | Yagi 1978; Zieba et al 2000 |

| | | | | | | |
|---|---|---|---|---|---|---|
| *Homo sapiens* | Tumor tissue and lung parenchyma | Schiff Base (SB) measurements as in Buege and Aust 1978 | Adenocarcinoma: 7.71 ±3.02 U SB (tumor tissue)<br>Small cell lung cancer: 4.63 ±2.28 U SB<br>Squamous cell lung cancer: 5.96 ±3.6 U SB<br>Large cell lung cancer: 8.76 ±2.91 U SB | Adenocarcenoma, small cell lung cancer, squamous cell lung cancer, and large cell lung cancer | Normalized to sample volume | Buege and Aust 1978; Zieba et al 2000 |

| | | | | | | |
|---|---|---|---|---|---|---|
| *Homo sapiens* | Tumor tissue and lung parenchyma | $H_2O_2$ measurement as in Ruch et al 1983 | Adenocarcinoma: 0.235 ±0.131 µmol $H_2O_2$ $L^{-1}$<br>Small cell lung cancer: 0.233 ±0.108 µmol $H_2O_2$ $L^{-1}$<br>Squamous cell lung cancer: 0.193 ±0.167 µmol $H_2O_2$ $L^{-1}$<br>Large cell lung cancer: 0.11 ±0.06 µmol $H_2O_2$ $L^{-1}$ | Adenocarcenoma, small cell lung cancer, squamous cell lung cancer, and large cell lung cancer | Normalized to sample volume | Ruch et al 1983; Zieba et al 2000 |
| *Homo sapiens* | Cervical tissue and tumor tissue | TBARS assay as in Esterbauer et al 1982 | 0.183 ±0.164 µmol $H_2O_2$ $L^{-1}$ (lung parenchyma) | Cervical cancer progression across stages I-IV | Normalized to sample mass | Esterbauer et al 1982; Ahmed et al 1999 |

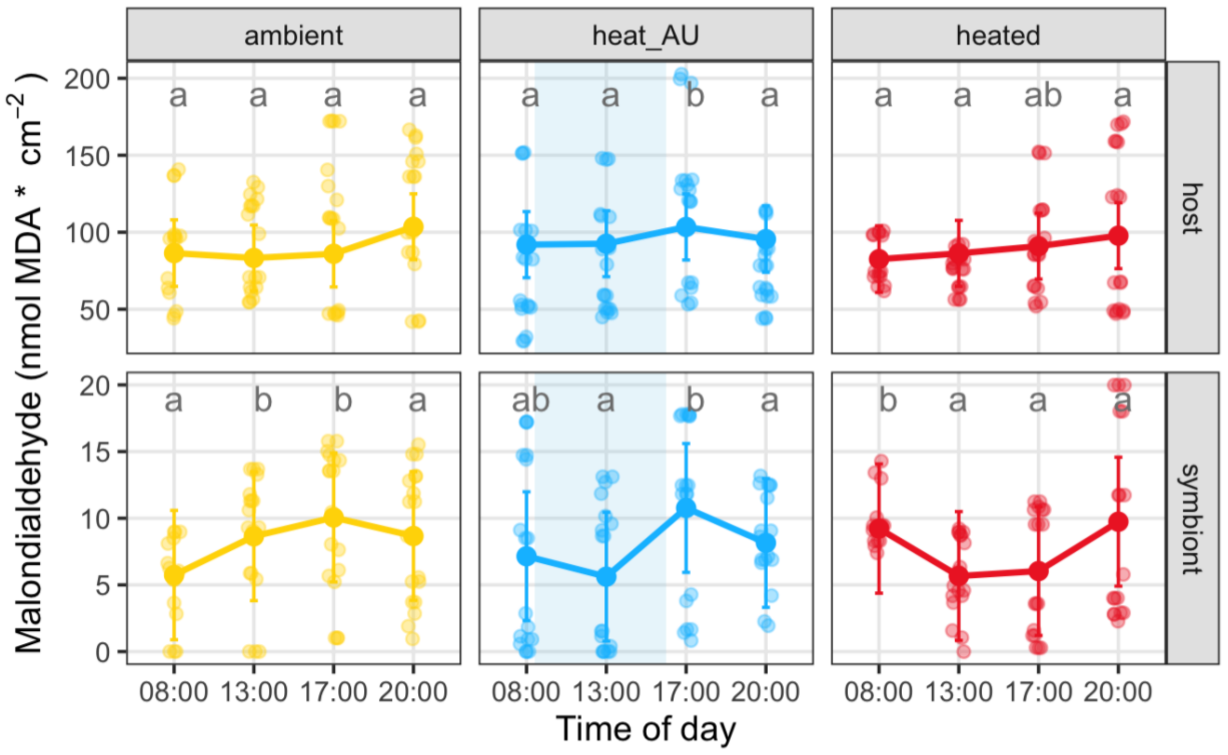


**Fig. S3** MDA concentration in *M. cavernosa* (nmol MDA mg$^{-1}$ protein) after exposure to ambient, heat_AU (artificial upwelling for 8h starting at 8:00 am), and heated treatments in the host and symbiont fractions, normalized to coral surface area. Opaque circles represent the mean MDA concentration, and partially transparent circles represent individual data points and replicates. Error bars depict standard error. The light blue shaded area indicates the cold water pumping period in the AU treatment. Letters indicate significant differences in pairwise, post hoc comparisons for treatments within timepoints

**Table S4.** MDA standard curves showing contradicting slope and $R^2$ values in Assays 1-5 on *M. cavernosa* samples.

| **Assay** | **Standard Curve Equation** | **Slope** | **$R^2$** | **Times standard stock was opened** |
|---|---|---|---|---|
| 1 | $A_{532} = 3.3389 \times [MDA]$ | 3.3389 | 0.9978 | 1 |
| 2 | $A_{532} = 3.2488 \times [MDA]$ | 3.2488 | 0.9992 | 1 |
| 3 | $A_{532} = 4.6552 \times [MDA]$ | 4.6552 | 0.9996 | 2 |
| 4 | $A_{532} = 16.591 \times [MDA]$ | 16.591 | 0.9957 | 3 |
| 5 | $A_{532} = 10.907 \times [MDA]$ | 10.907 | 0.9326 | 4 |
